# Preparation of DPPC liposomes using probe-tip sonication: investigating intrinsic factors affecting temperature phase transitions

**DOI:** 10.1101/742437

**Authors:** Monica D. Rieth, Andrew Lozano

## Abstract

Liposomes are an important tool and have gained much attention for their promise as an effective means of delivering small therapeutic compounds to targeted sites. In an effort to establish an effective method to produce liposomes from the lipid, dipalmitoyl-phosphatidylcholine or DPPC, we have found important aspects that must be taken into consideration. Here, we used probe-tip sonication to prepare liposomes on a batch scale. During this process we uncovered interesting steps in their preparation that altered the thermodynamic properties and phase transitions of the resulting liposome mixtures. Using differential scanning calorimetry to assess this we found that increasing the sonication time had the most dramatic effect on our sample, producing almost an entirely separate phase transition relative to the main phase transition. This result is consistent with reports from the current literature. We also highlight a smaller transition, which we attribute to traces of unincorporated lipid that seems to gradually disappear as the total lipid concentration decreases. Overall, sonication is an effective means of producing liposomes, but we cannot assert this method is optimal in producing them with precise physical properties. Here we highlight the physical effects at play during this process.

## Introduction

Liposomes are bilayered nanostructures typically comprised of phospholipids that can have variable properties, which ultimately influence the larger nanostructures they make up. Small peptides, nucleic acids and polymer-based materials can also be doped in to create liposomes with unique features making them adaptable for specific applications like drug delivery or other uses that require them to be stable.^1–5^ Other factors like pH, temperature and ionic strength can also play a role in affecting their inherent stability.^6–10^ They are also used as an effective tool in molecular biology to facilitate organism transformation and transfection with foreign DNA or RNA.^11–14^

In solution, methods like extrusion, sonication, and rapid ethanol injection have been used to successfully prepare liposomes.^15–17^ They can also be prepared from thin films and other synthetic supports.^18,19^ The application of each technique can influence the final physical properties including size (diameter), lamellarity (single bilayer or multi-bilayer), polydispersity (range of sizes), and zeta potential (surface charge) to name a few.^5,17,20,21^ In this study we used differential scanning calorimetry, DSC, as a means of assessing the thermal stability and phase transitions of liposomes prepared from a single lipid to help establish the parameters that are critical for their reproducibility by probe-tip sonication. DSC has previously been reported as a useful tool in studies assessing liposome stability. We chose this method because it is accepted as a means of evaluating liposome stability. We investigated three parameters in the development toward establishing a robust method of liposome preparation, which will later be employed for downstream applications. Upon investigating these parameters, we found that the quantity of lipid used for sample preparation and the total sonication time significantly affected the transition state temperature and overall sample stability, which is defined herein by a decrease in the melting temperature transition state compared to a defined standard and previously reported values in the literature. Our analysis stems from the creation and optimization of a consistent and reliable method to prepare pure DPPC liposomes, 1,2-dipalmitoyl-*sn*-glycero-3-phosphocholine, using probe-tip sonication. We chose to use DPPC because it is a relatively well-characterized lipid to which we could readily compare our results. The over-arching aim of this work is to use the insights generated from this investigation to reliably produce liposomes for the capture of small molecule compounds.

## Materials and Methods

### Sample preparation and DSC measurements

The saturated lipid, 1,2-dipalmitoyl-sn-glycero-3-phosphocholine (DPPC), was purchased from Avanti Polar Lipids (cat.# 850355, Alabaster, AL) and used without further purification. DPPC was weighed out to a mass of 2.0, 5.0, 10.0, and 25.0 mg. The lipids were transferred to a 2.0 mL glass vial to which 1 mL of buffer (20 mM HEPES pH 7.4, 100 mM NaCl) was added. The final volume did not exceed 1.0 mL. Samples were vortexed briefly to mix which resulted in a milky white suspension containing larger white particulates. To generate liposomes, the mixtures were sonicated using a probe-tip sonic dismembrator equipped with a microtip adapter (Fisher Scientific, Hampton, NH) set to 20% duty cycle (relative pulse intensity) with a pulse length time of 2 seconds and a rest period of 5 seconds for a total of 4 minutes. This cycle was carried out a total of three times on each sample to prevent excessive heating. The liposome solutions were then transferred to a 2 mL Eppendorf tube and centrifuged using a microcentrifuge, (Eppendorf, Model 5424) at 10,000 rpm for 3 minutes to remove titanium particles introduced during sonication. The supernatant was then transferred to another 2 mL Eppendorf tube. Samples were stored overnight at 4 °C and DSC studies were carried out the following day. Samples were not stored longer than 16-20 hours in advance of the DSC studies to preserve sample integrity and minimize liposome degradation. Stored liposome samples were removed from the refrigerator and left to equilibrate at room temperature for at least one hour. The samples were centrifuged at 10,000 rpm for an additional 3 minutes and the supernatant was carefully transferred to a clean 2.0 mL Eppendorf tube. All samples and buffer were degassed for approximately 30 minutes and the buffer was filtered prior to sample preparation using a 0.2 µm syringe filter.

DSC measurements were carried out on a VP-DSC high sensitivity scanning calorimeter (MicroCal, Northampton, MA, USA). All samples were scanned at a rate of 60 °C / hr from 20 °C to 70 °C. Samples were pre-equilibrated for five to ten minutes at 20 °C (room temperature) prior to the initial scan. To avoid the possibility of irreversible degradation, one scan per sample was obtained and replicates were carried out on freshly prepared samples.

### Changes in DSC temperature scan rates

Briefly, two samples were prepared at 5.0 mg / mL and each was sonicated using the probe-tip sonicator set to 20% duty cycle with a pulse length time of 2 seconds and a rest period of 5 seconds for a total of 4 minutes. This was repeated a total of three times for the two samples for a total of 12 minutes. The samples were centrifuged and stored overnight at 4 °C. DSC studies were carried out the following day.

DSC measurements were carried out as described above on each sample. The samples were scanned from 20 °C to 70 °C at a rate of 30 °C / hr and 60 °C / hr. Samples were pre-equilibrated for five to ten minutes at 20 °C prior to the initial scan.

### Varying the sonication time during liposome preparation

DPPC liposomes were prepared at a total lipid composition of 25.0 mg / mL. Five samples were weighed out to a mass of 25.0 mg. The lipids were transferred to a 2.0 mL glass vial and 1 mL of buffer was added. To generate liposomes, the mixtures were sonicated with the probe-tip sonicator for 6, 12, 20, 28, and 36 minutes set to 20% duty cycle with a pulse length time of 2 seconds and a rest period 5 seconds for a total of 4 minutes. This was repeated one and a half times for the 6-minute sample, three times for the 12-minute sample, five times for the 20-minute sample, seven times for the 28-minute sample, and nine times for the 36-minute sample. To avoid excessive heating the samples were rotated through one sonication cycle after 4 minutes. A 14-minute rest was introduced in between the last two cycles for the 36-minute sample to avoid excessive heating. The DPPC liposome solutions were then transferred to a 2 mL Eppendorf tube and centrifuged using a microcentrifuge (Eppendorf, Model 5424) at 10,000 rpm for 3 minutes to remove titanium particles introduced during sonication. The supernatant was transferred to a new 2 mL Eppendorf tube and samples were stored overnight at 4 °C. DSC measurements were carried out the following day. Samples were not stored longer than 16-20 hours prior to carrying out the DSC measurements to preserve sample integrity and minimize liposome degradation. The stored liposome samples were removed from the refrigerator, left to equilibrate at room temperature for at least one hour, centrifuged at 10,000 rpm for 3 minutes and degassed along with buffer for 30 minutes.

DSC measurements were carried out as described with a scanning temperature rate of 60 °C / hr beginning at 20 °C and ending at 70 °C. Samples were pre-equilibrated for five to ten minutes at 20 °C prior to the initial scan.

After each measurement the raw tabulated data was imported to KaleidaGraph version 4.5 (Synergy, Reading, PA) and plotted. Baseline subtraction was carried out manually prior to generating thermograms and extracting the thermodynamic properties. The main phase transition temperature (T_m_) was extracted and the peak morphology in each thermogram was analyzed for changes in the main phase transition temperature. Peaks arising from additional phase transitions were reported and summarized along with their thermodynamic parameters.

## Results

DPPC liposomes were prepared using probe-tip sonication, one of several methods that is commonly used. Liposome samples were prepared at 2.0, 5.0, 10.0, 25.0 mg/mL and DSC measurements were carried out. In all of the samples, a major temperature transition peak appears at approximately 41.0 °C, which is consistent with what has been previously reported for DPPC liposomes (Figure 1).^22^ A secondary transition peak begins to appear immediately before the main peak close to 38.0 °C. This secondary peak becomes increasingly noticeable as the lipid concentration increases above 5 mg / mL (Figure 1), although, when we lower the rate of the temperature scan we see an enhanced resolution of this secondary transition (Figure 2A).

**Figure 1.**
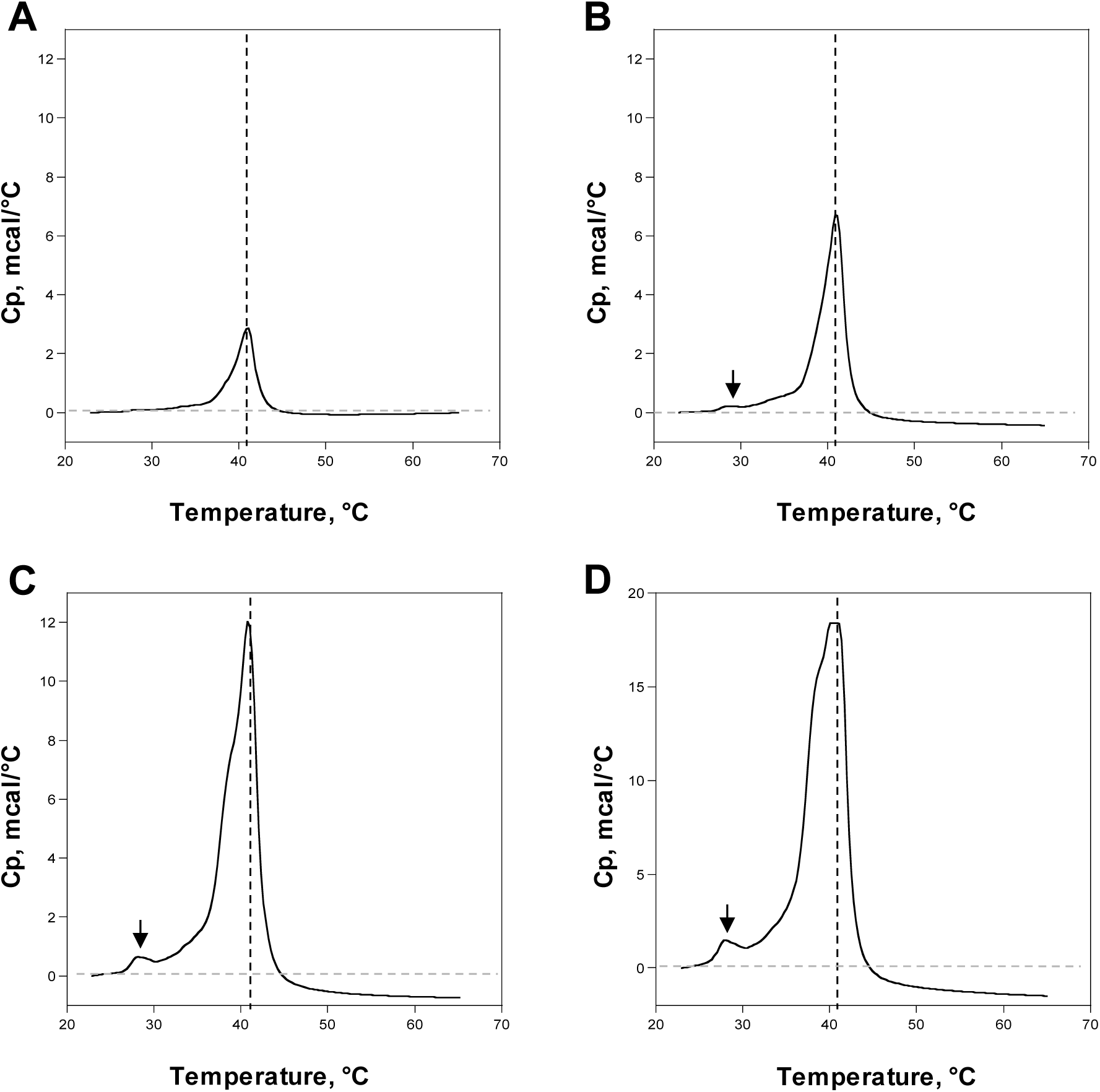
DSC thermograms of DPPC liposome samples prepared with A) 2.0 mg / mL, B) 5.0 mg / mL, C) 10.0 mg / mL, and D) 25.0 mg / mL. The curves have been normalized to zero and the major peak at approximately 41.0 °C is denoted by the dotted line. The small peak at approximately 27.0 °C denoted by the black arrow can be attributed to unincorporated lipid. All scans were performed at 60 °C / hr.

**Figure 2.**
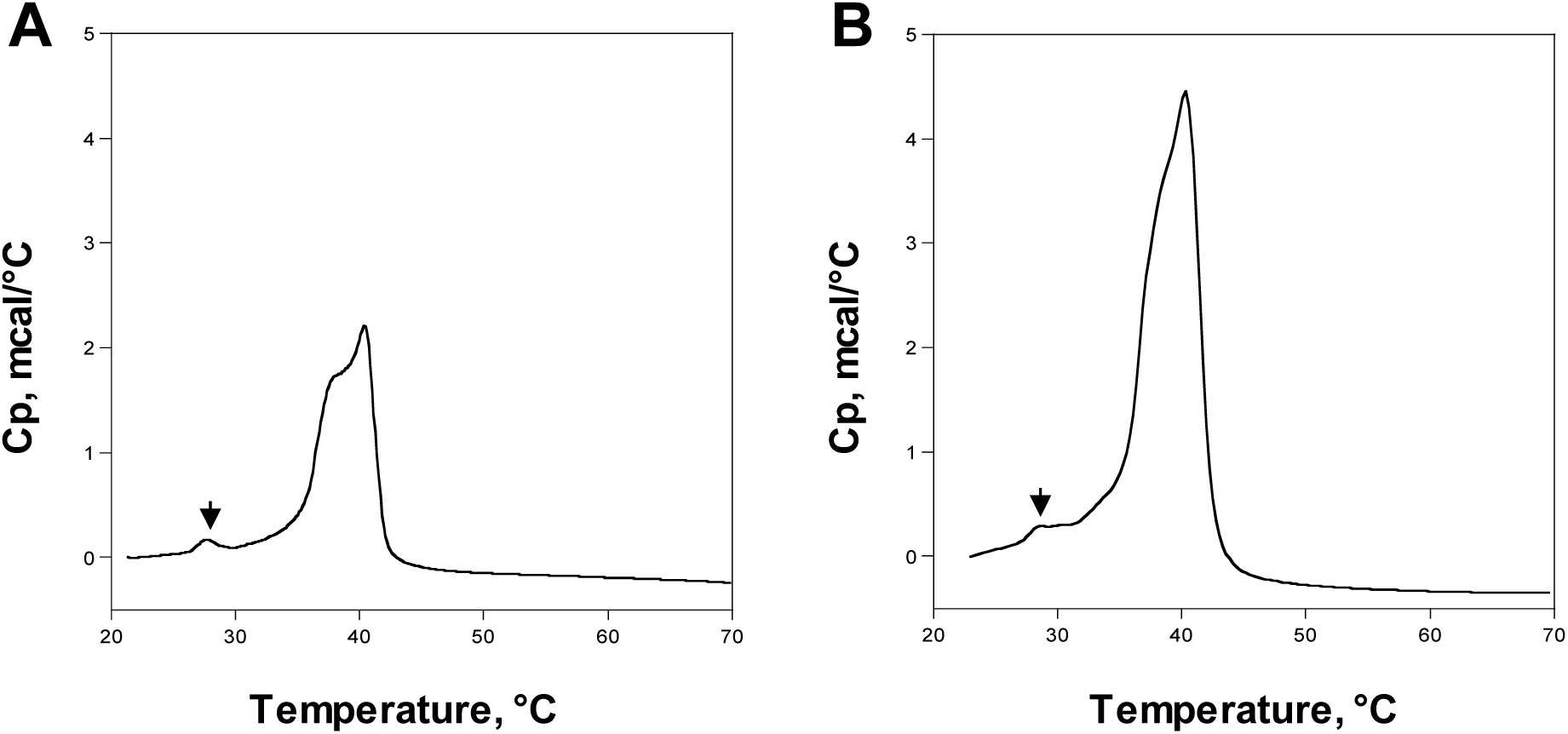
DSC thermogram of DPPC liposome samples prepared at 5.0 mg / mL. A) scanned at 30 °C / hr, B) scanned at 60 °C / hr. A subtle shoulder is present in both samples but becomes more apparent when the temperature scan rate is decreased by half.

We attribute this secondary temperature transition to the formation of another type of liposome population altogether, though, the appearance of this peak has been previously reported as a “pre-transition” peak prior to the main phase transition. This behavior has also been previously described for polymorphic mixtures.^23,24^ The secondary peak (Table 1 - minor peak) became more readily apparent when the temperature was increased more gradually at 30 °C / hr compared to 60 °C / hr (Figure 2).

**Table 1.**
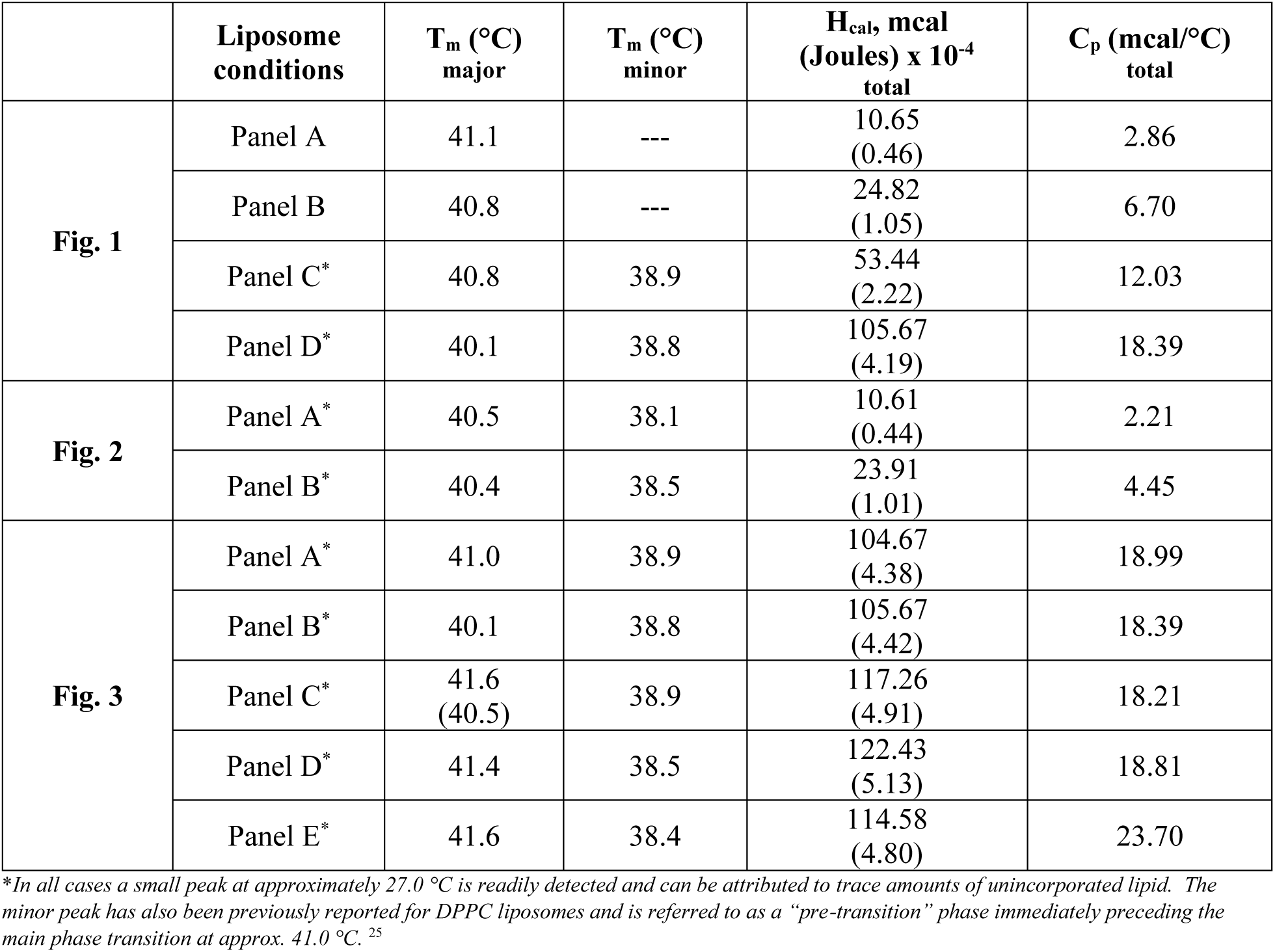
Summarized thermodynamic data extracted from DSC thermograms.

|  | **Liposome conditions** | **T<sub>m</sub> (°C) major** | **T<sub>m</sub> (°C) minor** | **H<sub>cal</sub>, mcal (Joules) x 10<sup>-4</sup> total** | **C<sub>p</sub> (mcal/°C) total** |
| --- | --- | --- | --- | --- | --- |
| **Fig. 1** | Panel A | 41.1 | --- | 10.65<br>(0.46) | 2.86 |
|  | Panel B | 40.8 | --- | 24.82<br>(1.05) | 6.70 |
|  | Panel C* | 40.8 | 38.9 | 53.44<br>(2.22) | 12.03 |
|  | Panel D* | 40.1 | 38.8 | 105.67<br>(4.19) | 18.39 |
| **Fig. 2** | Panel A* | 40.5 | 38.1 | 10.61<br>(0.44) | 2.21 |
|  | Panel B* | 40.4 | 38.5 | 23.91<br>(1.01) | 4.45 |
| **Fig. 3** | Panel A* | 41.0 | 38.9 | 104.67<br>(4.38) | 18.99 |
|  | Panel B* | 40.1 | 38.8 | 105.67<br>(4.42) | 18.39 |
|  | Panel C* | 41.6<br>(40.5) | 38.9 | 117.26<br>(4.91) | 18.21 |
|  | Panel D* | 41.4 | 38.5 | 122.43<br>(5.13) | 18.81 |
|  | Panel E* | 41.6 | 38.4 | 114.58<br>(4.80) | 23.70 |

\*In all cases a small peak at approximately 27.0 °C is readily detected and can be attributed to trace amounts of unincorporated lipid. The minor peak has also been previously reported for DPPC liposomes and is referred to as a “pre-transition” phase immediately preceding the main phase transition at approx. 41.0 °C.<sup>25</sup>

In nearly all of the samples a small peak is also readily apparent at 27.0 °C (denoted by the small black arrow), but is only slightly detected at concentrations below 5 mg / mL (Figure 1). We attribute this to unincorporated lipid that could not be removed by centrifuging after sonication. Next, we altered the total sonication time for five samples. The sonication time ranged from 6 to 36 minutes. Interestingly, the secondary peak that emerged from the previous experiment becomes remarkably more pronounced when the sonication time was increased to 36 minutes and we believe this may be attributed to a phase transition arising from a distinct new liposome species that was not present before (Figure 3). This secondary peak, again, appears between T_m_ = 38.4-38.9 °C, which is similar to what we see in Figures 1 and 2. The small peak between 27.0-29.0 °C is still present in each of the samples, which were all prepared with 25.0 mg / mL DPPC and scanned at a rate of 60 °C / hr.

**Figure 3.**
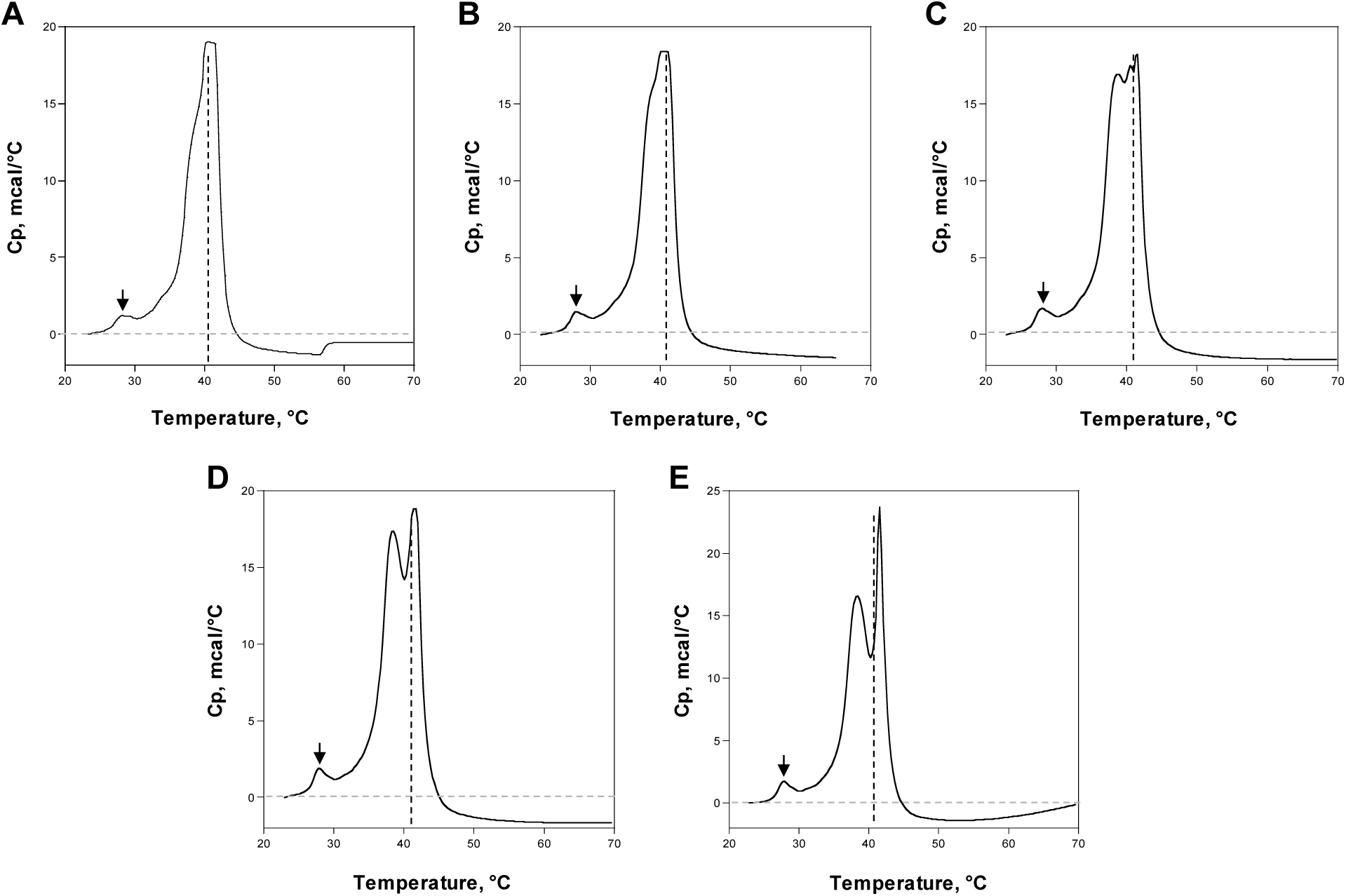
DSC thermogram of DPPC liposome prepared at different total sonication times, A) 6 minutes, B) 12 minutes, C) 20 minutes, D) 28 minutes, E) 36 minutes. The transition temperature introduced by unincorporated lipid is denoted by the small arrow. All scans were performed at 60 °C / hr with 25.0 mg / mL DPPC.

The thermodynamic data from each DSC thermogram is summarized in Table 1 and shows the melting temperature at each major phase transition, T_m_, for the lipid species in each thermogram. The heat capacity, C_p_, along with total calorimetric enthalpy, H_cal_, is also reported. Molar heat capacity and enthalpy were omitted due to the inherent inaccuracy of this prediction based on observations of reported lipid loss after sonication. Results are summarized and enthalpy is reported in both millicalories and in Joules. In all samples the major temperature phase transition occurred between 40.1 and 40.8 °C, which is consistent with what has been previously reported.

## Discussion

In the preparation of liposome mixtures, thermal stability is often an important parameter to consider, especially when attempting to prepare samples for downstream applications. The methods used to prepare liposomes can often have important consequences and ultimately must be considered when attempting to establish reproducible methods for producing a well-defined, homogeneous population with specified properties.^16^ Differential scanning calorimetry, DSC, is an accepted means of assessing the thermal stability and we chose to implement this method in the current study.^23,26,27^ Based on the method of probe-tip sonication, we further investigated parameters to determine which factors had the greatest influence on our liposome preparations. Though at first varying these parameters seemed trivial, the results from the analysis suggest otherwise and show these factors give rise to additional phase transitions, which we might not have otherwise predicted. Though there are other ways to prepare liposomes, this method is currently under development toward an established, reliable method from which additional experiments can be carried out. Although probe-tip sonication is relatively straightforward it is not the optimal method for producing liposomes with uniform properties. For example, lamellarity (unilamellar vs. multilamellar) and size (nm) can be drastically affected by the method of preparation.^18,20,28^ To produce liposomes with uniform size and lamellarity (single-bilayer or unilamellar) extrusion is often a favorable choice, however, the over-extending goals of our work are to explore the utility of liposome mixtures in capturing environmental compounds. To that end, producing liposomes with ultra-precise characteristics becomes less important compared to establishing the most ideal method based on our goals. While preparing our samples, we discovered interesting parameters that affected the thermal stability and observed changes in the phase transition of pure DPPC liposomes. To capture additional, though subtle, transition states it was more effective to increase the temperature gradually at 30 °C / hr for each DSC measurement, which increased peak resolution in the thermograms (Figure 2). Variations in sonication time were implemented to determine if this affected the overall thermal stability of the mixture and was motivated by our observation that there was substantial lipid loss after sonication. To ensure complete incorporation of all lipid material we tested variations in sonication time. Our standard practice describes a total sonication time of 12 minutes with a rest period introduced after 4 minutes to allow the sample to sufficiently cool and prevent overheating. Sample overheating can lead to lipid degradation which can affect the final properties of the liposomes, so it was important to carefully monitor samples for increases in temperature while extending the sonication time.^29^ Conversely, samples could not be chilled as a measure to prevent overheating during sonication else we introduce the risk of creating conditions whereby DPPC preferentially assumes a solid crystalline-like state. This can hinder liposome formation.^30^ We found there was noticeably less residual, unincorporated lipid that had accumulated at the bottom of the Eppendorf tube after centrifugation when the sonication time was increased. After 36 minutes there were virtually no traces of detectable lipid present at the bottom of the tube, however in all cases titanium particles were still present. Regardless of the presence or absence of a visible white pellet after sonication, traces of unincorporated lipid could still be detected in the DSC, which are highlighted in each figure by a small black arrow. This appears to vary proportionately with the amount of lipid initially present in the sample.

This work, though preliminary, reveals important aspects of liposome preparation that should be addressed when attempting to establish a reproducible method. Beyond the practical considerations of this work further studies that probe the biophysical behavior of lipids prepared this way may uncover interesting phenomena that we never could have predicted similar to the findings herein.

## Conclusions

We demonstrated in this brief analysis that small changes in the preparation of DPPC liposomes using probe-tip sonication can affect the thermal transition profile. Differential scanning calorimetry has been used as a tool to assess liposome stability and measure temperature transition states and here we employ this as means to assess important parameters that should be considered when attempting to establish robust and reproducible preparatory methods specifically using probe-tip sonication.^23,26^ The appearance of additional peaks relative to the main transition peak at approximately 41.0 °C suggests the presence of additional liposomal species or the generation of a polymorphic mixture when lipid concentrations are increased and sonication times are extended, which has been previously reported for pure DPPC liposomes.^23^

## Acknowledgements

The authors acknowledge the support of the Chemistry department.

